# Panama: A tool for ion channel biophysics simulation

**DOI:** 10.1101/202051

**Authors:** Anuj Guruacharya

**Affiliations:** Department of Biology, University of Oklahoma, Norman, OK, 4058898692

## Abstract

I have created an online tool and an R library that simulates biophysics of voltage-gated ion channels. It is made publicly available as an R library called Panama at github.com/anuj2054/panama and as a web app at neuronsimulator.com. A need for such a tool was observed after surveying available software packages. I found that the available packages are either not robust enough to simulate multiple ion channels, too complicated, usable only as desktop software, not optimized for mobile devices, not interactive, lacking intuitive graphical controls, or not appropriate for undergraduate education. My app simulates the physiology of 11 different channels - voltage-gated sodium, potassium, and chloride channels; channels causing A-current, M-current, and After-HyperPolarization (AHP) current; calcium-activated potassium channels; low threshold T type calcium channels and high threshold L type calcium channels; leak sodium and leak potassium channels. It can simulate these channels under both current clamp and voltage clamp conditions. As we change the input values on the app, the output can be instantaneously visualized on the web browser and downloaded as a data table to be further analyzed in a spreadsheet program. The app is a first of its kind, mobile-friendly and touch-screen-friendly online tool that can be used to teach undergraduate neuroscience classes. It can also be used by researchers on their local computers as part of an R library. It has intuitive touch-optimized controls, instantaneous graphical output, and yet is pedagogically robust for education and casual research purposes.

Neuroscience education, ion channel biophysics, Hodgkin-Huxley simulation, web app for neuroscience

## 1 Introduction

The Hodgkin-Huxley (Hodgkin et.al., 1952) model is one of the fundamental neuronal models. Its mathematical form is a set of differential equations that students in a neuroscience class are taught before moving on to more complex models. Computational simulations using this model strengthen students’ concepts of action potentials and ion channels.

Existing simulation programs, such as NEURON (Hines and Carnevale, 1997) and GENESIS (Bower et al., 2007), serve as powerful tools for simulating the response of whole-cell or single-channel parameters to electrical or pharmacological stimuli. Although such software tools are free and could be used for educational or research purposes, they require substantial training and may not be suitable for casual use by researchers. Some effort has been directed towards making educational packages that demonstrate ion channel biophysics that are freely available. These are good tools to teach students about action potentials, ion channel currents and voltages. However, each of these tools has its own disadvantages. I highlight some of them in particular. HHsim ( http://www.cs.cmu.edu/dst/HHsim/) (Touretzky et al., 2003) requires the software to be downloaded and a matching version of Matlab installed on the desktop computer. Neurophysiology Virtual Lab (http://vlab.amrita.edu/?sub=3&brch=43) (Sridharan et al., 2016) requires a signup procedure and is not mobile-friendly. NeuroLab (http://sites.lafayette.edu/schettil/neurolab/) (Schettino, 2014) requires a special software environment called Netlogo. Others such as Phet (https://phet.colorado.edu/en/simulation/neuron>) are cartoon reconstructions of ion channel physiology with restricted features. A webapp located at http://myselph.de/hodgkinHuxley.html serves a sufficient purpose in it being an online simulator, but is not touch-optimized, and does not have enough adjustable input controls to be adequate for an undergraduate neurophysiology class. Nerve (http://nerve.bsd.uchicago.edu/) is not touch-or mobile-optimized while others, such as the programs available at http://www.eng.utoledo.edu/smolitor/download.htm (Molitar et al., 2006), are Matlab packages. Any software package that is dependent on Matlab is not ideal for wide distribution because of the overwhelming cost of Matlab and the requirement to preinstall Matlab. Any software package dependent on Java in the browser is not ideal because of the unavailability of built-in Java support in some modern web browsers.

Thus, a need was felt to make a tool that had the following characteristics: 1) mobile-friendly, 2) touch-screen-friendly, 3) pedagogically adequate for an undergraduate neurophysiology class, 4) completely online, 5) not reliant on Matlab or Java software, 6) built with open sourced code, and 7) usable by researchers that want an intuitive way to change ion channel parameters and download the data. To our knowledge, no electrophysiology simulation tool exists that satisfied all these criteria. In this paper, a new webapp for simulating the biophysics of voltage-gated ion channels is described. It has been made publicly available at http://www.neuronsimulator.com and as a downloadable R package called Panama through Github. The design of my simulator overcomes the limitations of previous simulators and satisfies all the criteria listed previously. R (R Core Team, 2014), Shiny package (RStudio Inc., 2014), and Lattice package (Sarkar, 2008) were used to code the software. It has multiple input controls for both voltage clamp and current clamp conditions. It outputs the voltage, current, and conductance values as graphs for each ion channel.

## 2 Methods

### 2.1 Numerical design of the simulator

The 11 channels simulated in this app were voltage-gated sodium, potassium, and chloride channels; calcium-activated potassium channels (*KCa*); T-type calcium channels (*CaT*); L-type calcium channels (*CaL*), leak sodium (*NaLeak*), and leak potassium (*KLeak*) channels; A current channels; M current channels; and AHP current channels. Each channel was represented by its maximal conductance or permeability (*g*_*n*_ or *p*_*n*_ where n is the specific ion channel), its ionic current (*I*_*n*_), its reversal potential (*E*_*n*_) and its associated gating parameters. Total ionic current (*I*_*net*_) was modeled as the sum of all those individual Hodgkin-Huxley style ionic currents: *i*_*Na*_, *i*_*K*_, *i*_*Cl*_, *i*_*NaLeak*_, *i*_*KLeak*_,*i*_*A*_, *i*_*M*_, *i*_*KCa*_, *i*_*AHP*_, *i*_*KCa*_, *i*_*M*_, *i*_*CaT*_, *i*_*CaL*_. The models for these channels were modified from those used in the EOTN software (Huguenard et al.,1996).

Voltage or current across the membrane was held constant depending on the clamping conditions. For the current clamp case, *I*_*net*_ was held for the clamp duration at the applied current provided by the user; *V*_*net*_ was determined from Kirchoff’s current law by solving a differential equation given in Equation 1. The membrane capacitance per area represented by *C* in Equation 1 is input by the user and set to a default value of 0.01 nFarads.

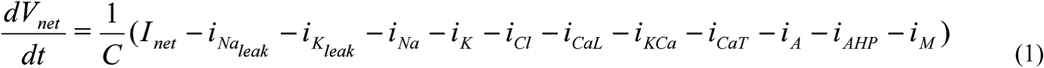

For the voltage clamp case, *V*_*net*_ was held for the clamp duration (set to a default of 50 ms) at the applied voltage provided by the user; *I*_*net*_ was determined from Kirchhoff’s current law as shown in Equation 2.

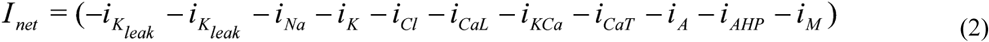

The default simulation time was set to 70 ms, with 10 ms being the pre-clamp and 10 ms being the post-clamp duration. This helped avoid data overflow issues. However the user can change the clamp duration, pre-clamp duration and post-clamp duration to increase the total simulation time to 1000 ms.

Equations 3 to 13 described the individual ionic currents for each channel used in the simulator. The electrochemical gradients driving the flow of ions were represented as voltage sources (*E*_*n*_) whose voltages were determined by Nernst equations that are dependent on the ratio of extra-to intra-cellular concentrations of the ionic species of interest. All gating variables, *m, n, h, m*_*KCa*_, *m1*_*A*_, *m2*_*A*_, *h1*_*A*_, *h2*_*A*_, *m*_*AHP*_, *m*_*M*_, *m*_*T*_, *m*_*L*_ were dimensionless numbers, ranging from 0 to 1. *m* denotes the probability of finding a channel in its open/permissive state and 1-*m* denotes the probability of it being closed or in a non-permissive state (current flow will be zero in this state). Other gating variables followed a similar representation. *InCa* and *OutCa* represented the concentration of calcium ions inside the cell and outside the cell in mM. The constant values used in Equations 5 to 16 which cannot be changed by the user are the temperature (*T*) set to be a room temperature of 293.15 Kelvins, gas constant (R) set to 8314 Joules/mole/kelvin, and the Faraday constant (*F*) set to 96485 Ampere-seconds/mole.

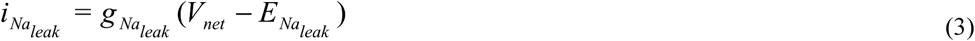

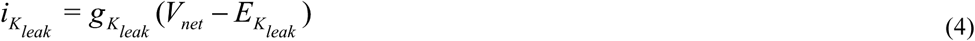

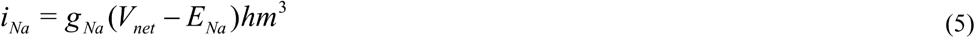

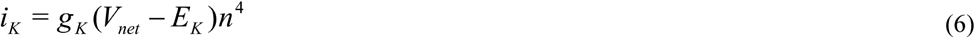

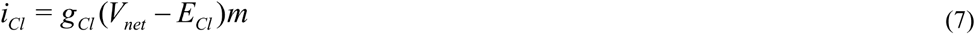

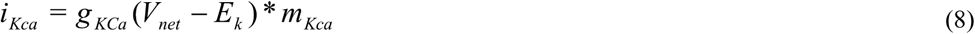

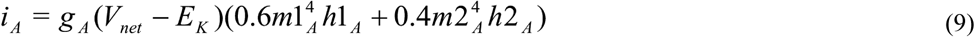

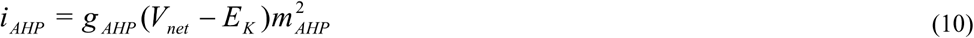

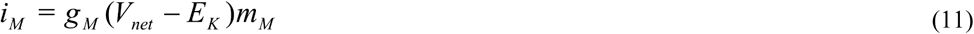

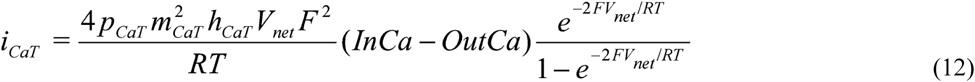

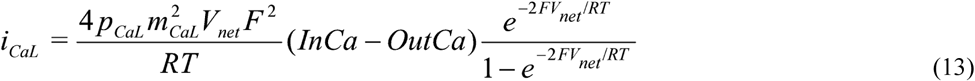

Differential equations in Equations 14 to 27 were used to model the activation and deactivation of the gating variables. The differential equations were discretized using an explicit forward Euler algorithm with a default time step of 0.025 ms. The time step can be changed by the user, however it is not recommended to turn it lower than the default value since it can slow down the simulation. The equations were assumed to be initial value problems and solved for each value of voltage across the membrane. So these differential equations were solved with different values of *V*_*net*_ across the membrane.

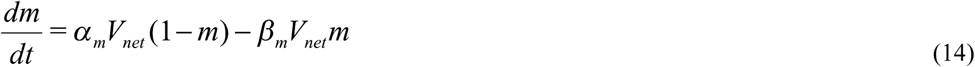

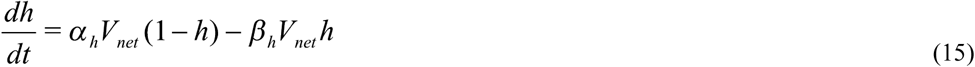

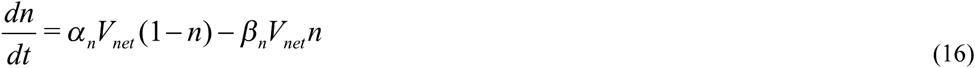

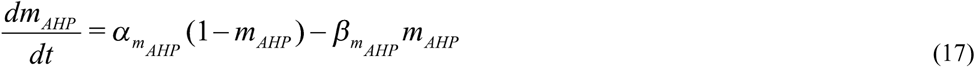

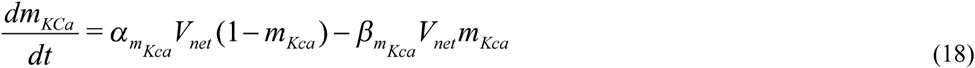

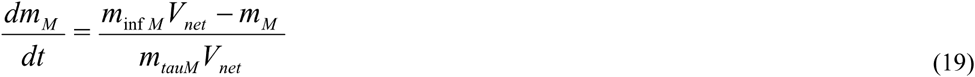

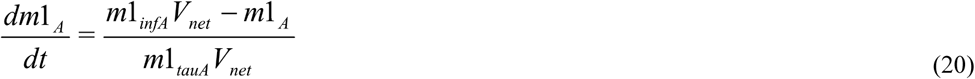

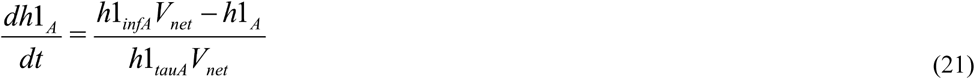

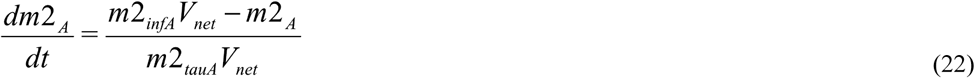

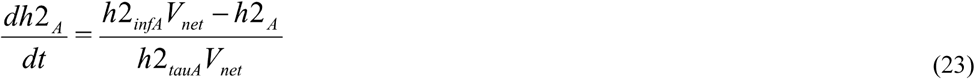

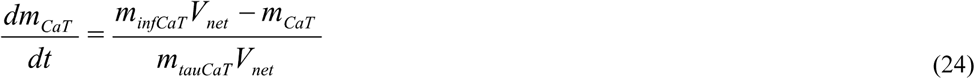

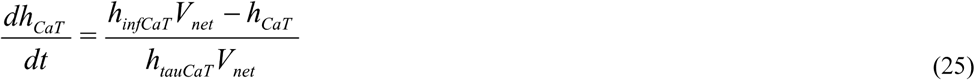

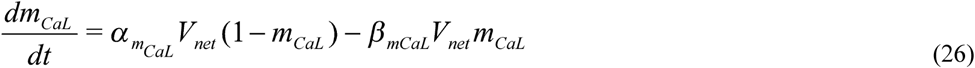

In all models, voltage was measured in *mVolt*, current in *nAmp*, time in *msec*, conductance in *μSiemens*, and capacitance in *nFarads*. The model parameters for sodium channels were derived from data of Huguenard et al. (1988). The potassium channel model used was of a general delayed rectifier. T-type calcium channel (*i*_*CaT*_) was modelled using the constant field equation. L-type calcium channel (*i*_*CaL*_) was also modelled using constant field equation as in *i*_*CaT*_, except that it was considered to not inactivate. *i*_*CaL*_ was based upon the data of Kay and Wong (1987) from isolated hippocampal pyramidal cells. Calcium activated potassim channel (*i*_*KCa*_) was modelled according to the procedure used by Yamada et al. (1989). The A current was modelled to inactivate with two time constants. The first component comprising of *m1*_*A*_ and *h1*_*A*_ contributes 60% to the total value of the gating variables. The second component comprising of *m2*_*A*_ and *h2*_*A*_ contributes 40% to the total value of the gating variables. The M current model was adapted from Adams et al. (1982). The AHP current was modelled according to the model in bullfrog sympathetic cells by Yamada et al. (1989).

### 2.2 Software design of the simulator

The webapp was created using the R programming language. After an initial survey of different languages and packages available in each language, the R language was chosen for its availability of Shiny and Lattice packages which are both excellent packages for web development and graphics development. R was also chosen because of its widespread use by biologists and undergraduates. The differential equations were coded into the script using only R, without using any external differential equation solver packages such as deSolve.

The Shiny package was used to serve the webpages. The Twitter bootstrap toolkit was used as the theme for user interface controls. Sliders from the bootstrap UI toolkit were used to make the input controls touch-friendly so that users do not have to type the values in a textbox.

The Lattice package was used to create the graphs that were embedded into the webpage. The output of the webapp is a set of voltage, current and conductance graphs for the channels. These can be visualized instantaneously while changing the input values on the app after pressing the update button, or they can be downloaded as CSV tables and analyzed in a spreadsheet software.

The code is open-sourced and deposited at http://www.github.com/anuj2054/panama. The front end of the software is coded in a file *ui.R* and the backend is coded in a file *server.R* as shown in Figure I. The app is hosted on a Shiny server located the the High Performance Computing Center facilities at Oklahoma State University. The computations for the equations all occur on the server’s side, so that there is no load on the user’s computer.

**Figure I:**
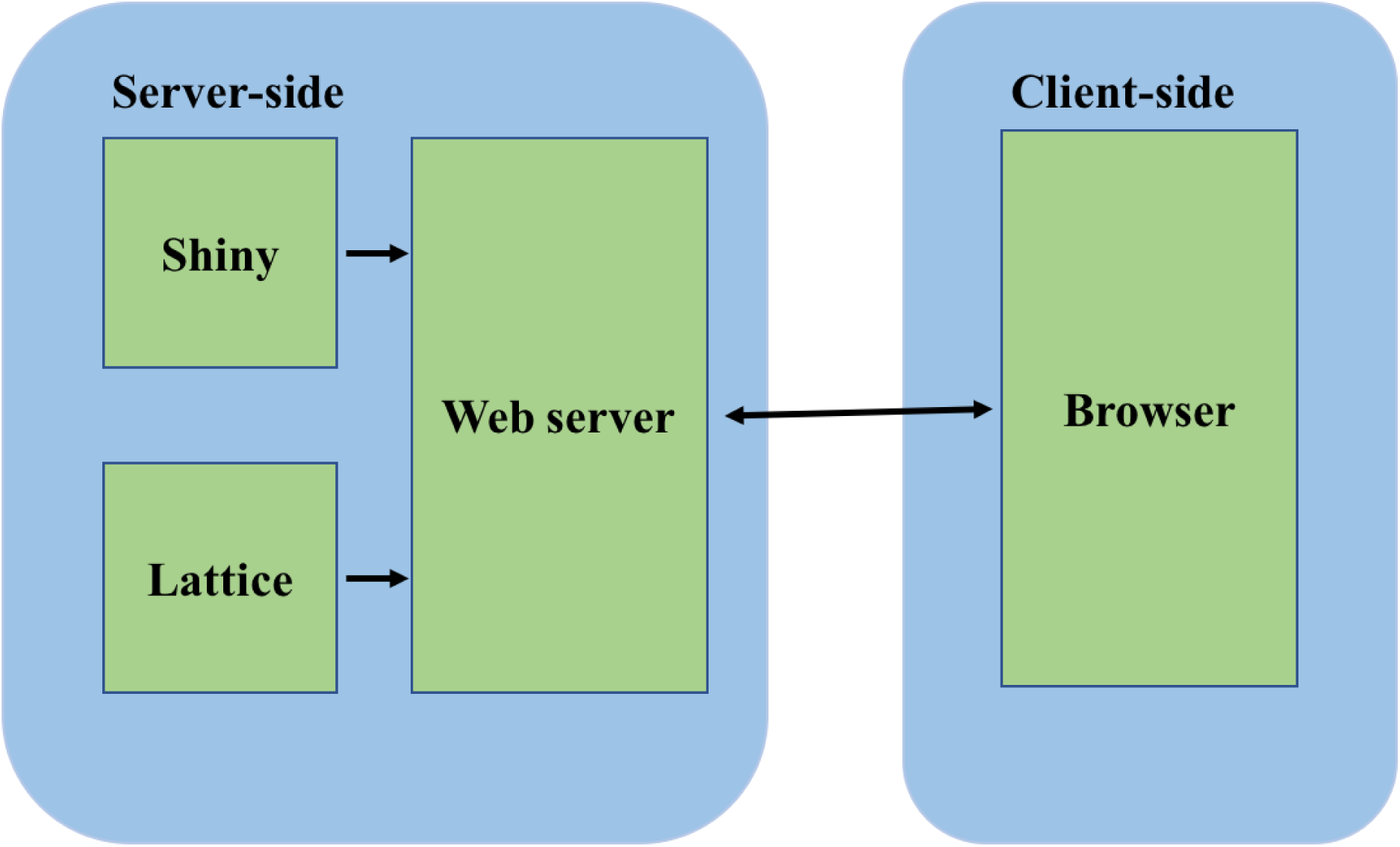
Software architecture of the simulator. The server side consists of the Shiny package and the Lattice package. The Shiny package is used for server side computations. The Lattice package is used for graphical display. The computations and the graphs are served to the client browser through the webserver built into the Shiny package.

## 3 Results and Discussion

There are three ways to access Panama. The first and the easiest way to access it is at http://www.neuronsimulator.com/. A second way that does not require a constant internet connection to work with the software is by using the command runUrl(‘https://github.com/anuj2054/panama/archive/master.zip’) on the R terminal. The command downloads the required files into an R folder and executes the software from the user’s local computer. A third way to access the app is by using the command shiny∷runGitHub(‘panama’, ‘anuj2054’) on the *R* terminal given that the user has preinstalled Shiny. In the second and third methods, once the required command is run and the code is automatically downloaded to the local computer, access to internet is not required anymore.

On a desktop web browser, the input controls appear as in Figure II. However, on a mobile device, the three columns of the input controls are merged into one column for easy scrolling. On a mobile device, the use of sliders eases the process of entering values for individual parameters of the model. However precision is not sacrificed. The user can change the input parameter values by hovering above the slider and using the keyboard to fine tune the exact number they want to three significant decimal places. The default values on the app can be reloaded by refreshing the web browser.

**Figure II:**
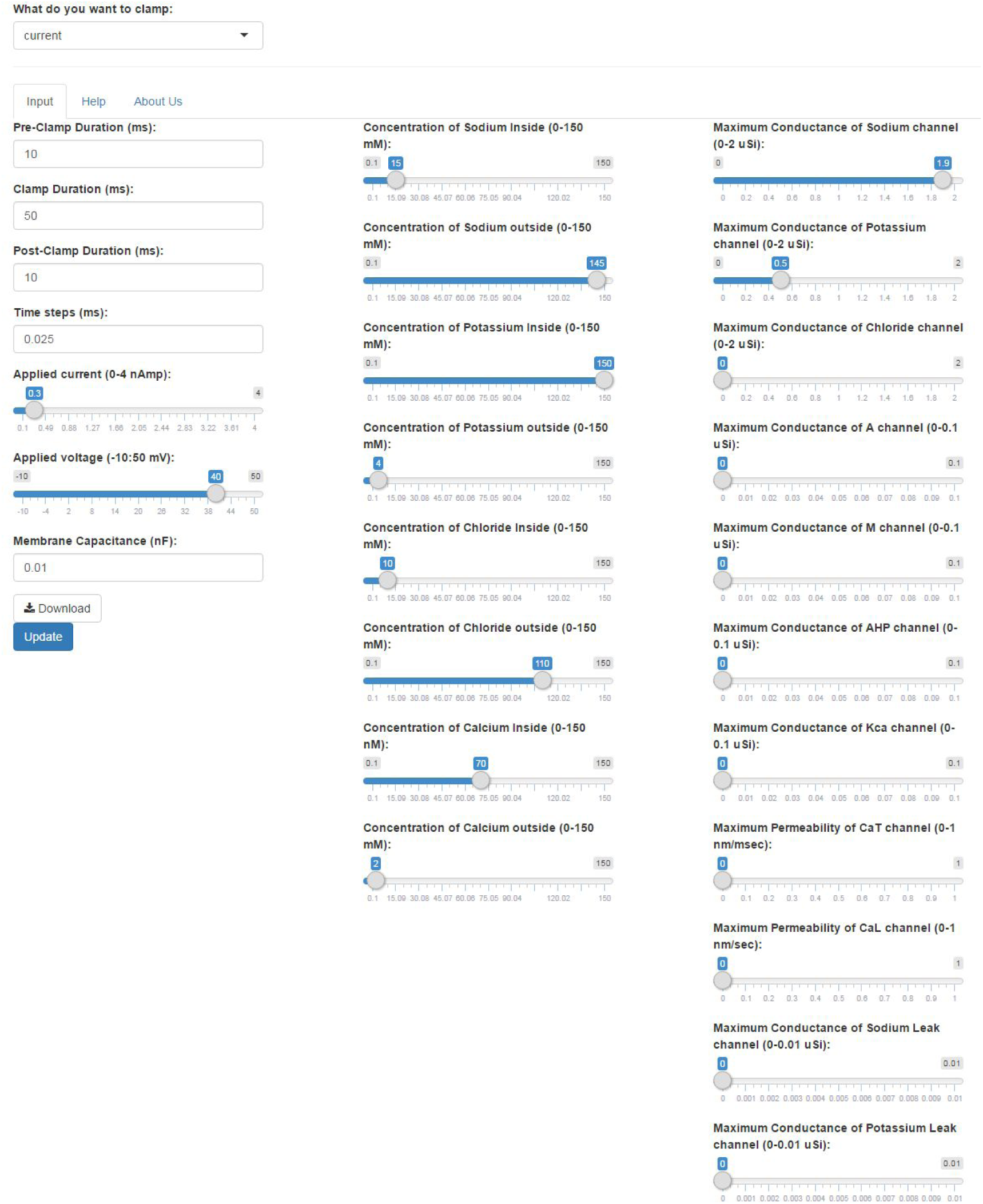
The input controls of the app on a desktop. The user first has to select the current clamp or voltage clamp conditions from the drop down menu. After selecting the clamping conditions, various parameters can be changed using the scrollbar. Pressing the *Update* button will update the output graphs depending on the values input. The user has to click on *Update* each time they change the parameters. Clicking on the *Download* button will download the time series voltage, current and conductances of the ion channels in a CSV format.

The app outputs conductance, voltage, and current data as both a graphical display and as downloadable CSV tables. The ability to download the output data as CSV tables enables the user to use their own spreadsheet software such as Excel to further analyze the data or embedd the graphs in their own documents. On a desktop web browser, the output graphs appear as shown in Figure III. Each of the lines inside the graphs are color coded and described with the name of the channel inside its respective colored rectangle. During the current clamp step, the current injection can be made more noticeable in the current graph by increasing the applied current and observing the steep red line that appears after the pre-clamp duration. The user can increase the clamp duration to see numerous action potentials that resemble neural spikes.

**Figure III:**
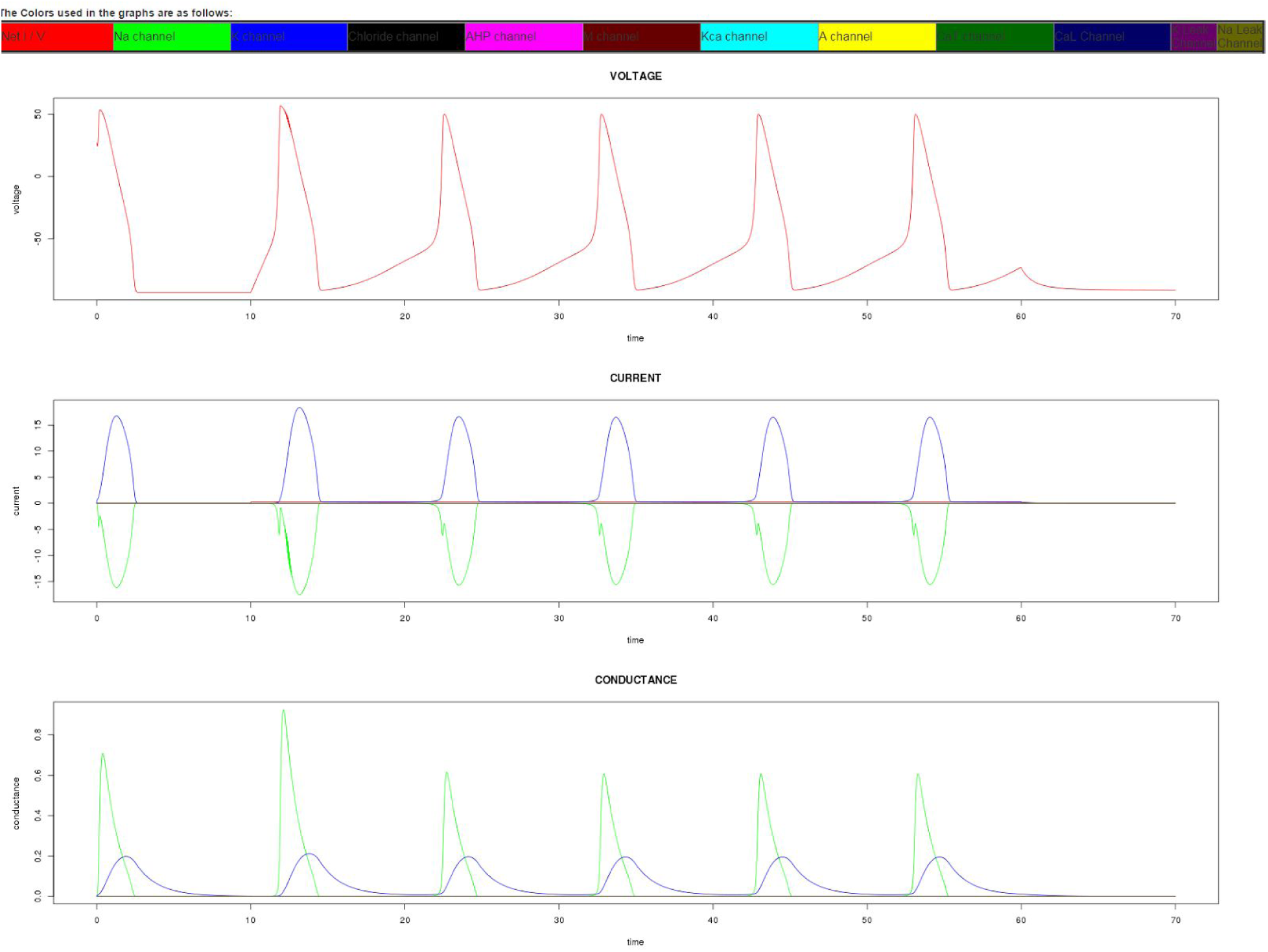
Output graphs of the app on a desktop. The simulator gives as output the time series representation of voltage, current, and conductance. The lines in the graph are color coded to represent different ion channels. In this particular graph, only sodium and potassium channels have been activated. The output graphs can also be replicated by downloading the CSV file and using a spreadsheet program to draw the graphs.

Users and students are encouraged to use the app in conjunction with a textbook about ion channels and their electrical properties. They are encouraged to change the default ion concentrations and channel conductances to those for different types of cells such as squid axon cells and observe its effect on the current and voltage of the cell. The default values used in the app are for a mammalian cell at room temperature (Lodish, 2008). During student interaction, it has been particularly helpful in pointing out to them the reasons for the particular changes they observe in the action potentials when different types of channels are activated. Changing the default capacitance and conductance values demonstrates to students how different conditions of a cell membrane affect the electrical properties of the cell. This practise gives students a hands on approach towards learning neurophysiology that they would otherwise only get from a textbook or from an expensive electrophysiology rig.

The numerical output of the simulator was tested against NEURON with similar parameters. Both programs returned equivalent results. The app was also tested under different operating systems (Windows, Android, iOS, Mac, and Linux) and under different browsers (Chrome, Firefox, and Internet Explorer). It was found to operate consistently across all platforms. The app has been well received by practicing and learning neurobiologists who have been exposed to its early versions. User behavior was tested on 3 volunteers. Based on their suggestions, the output graphs were color-coded and a simpler web address, neuronsimulator.com, was bought for the app. Future versions of the app will have phase space graphs to help users better understand membrane dynamics. It will also model synaptic currents where the chloride channels would play an important role. A better help section, tutorials for undergraduate students, and an even cleaner user interface is also being planned.

Panama is a first of its kind, a touch-friendly, mobile-friendly online tool that models electrophysiology of 11 different types of ion channels using Hodgkin-Huxley style differential equations. It requires no user training for installation and no infrastructure for download, making it suitable for casual research use or classroom environments. The webapp or the R package can be used by teachers to teach undergraduate neuroscience classes and also by researchers for casual use.

## Information Sharing Statement

The Panama software is available at www.neuronsimulator.com. Its associated scripts are available athttps://github.com/anuj2054/panama. R software is available at https://www.r-project.org/. Shiny Server is availableat https://www.rstudio.com/products/shiny/. Lattice software is available at https://cran.r-project.org/web/packages/lattice/index.html.

## Acknowledgments

This work was also facilitated utilizing the High Performance Computing Center facilities of Oklahoma State University (OSU) in Stillwater, Oklahoma. The author would like to thank Ari Berkowitz, Alex Bernard, Binita Rajbanshi, and Sharmistha Shayamal for their helpful comments on the manuscript. The author would also like to thank Jesse Schafer and Evan Linde for their support of computing facilities at OSU.

## Supplementary Materials

Equations 27 to 54 were used to model the rate constants used and the steady state values of the gating variables in Equations 14 to 26. These equations are dependent on the membrane voltage. For the initial value of *V*_*net*_ and *I*_*net*_, default values of 40 mVolt and 0.3 nAmp were used.

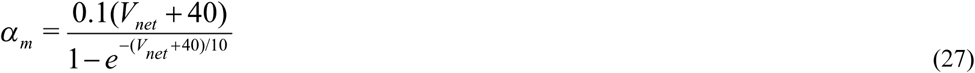

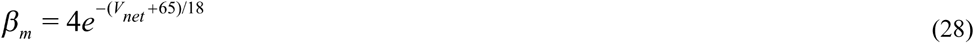

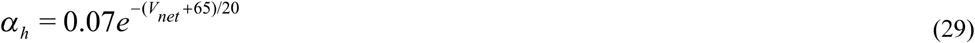

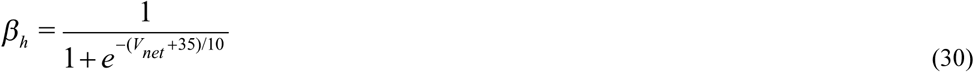

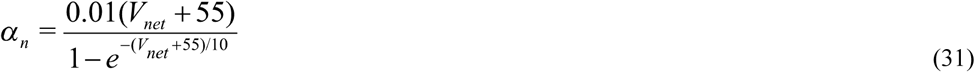

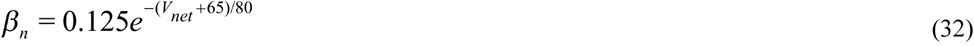

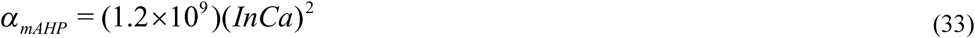

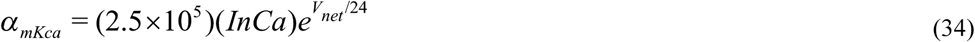

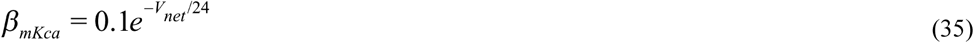

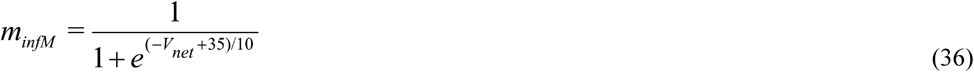

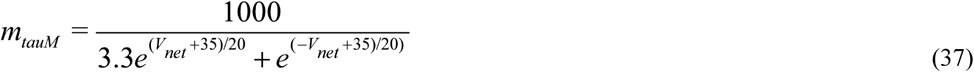

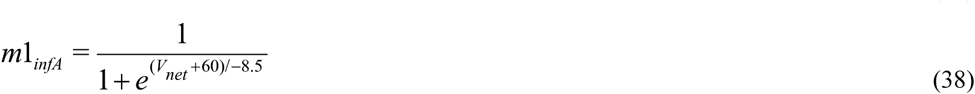

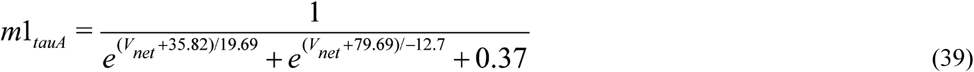

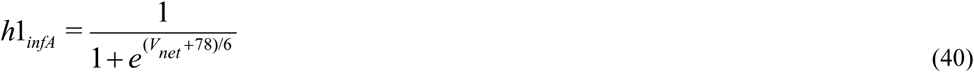

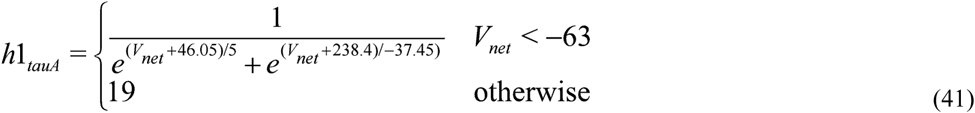

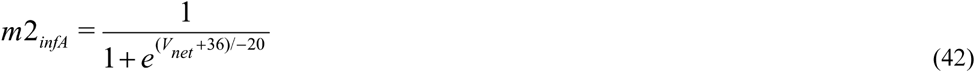

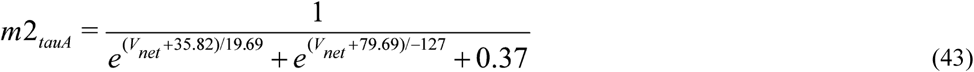

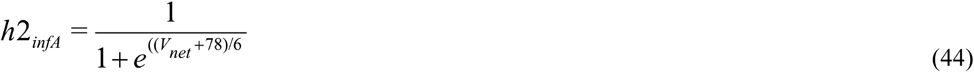

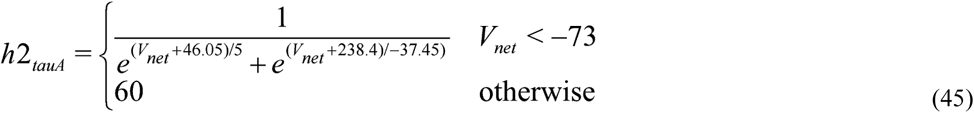

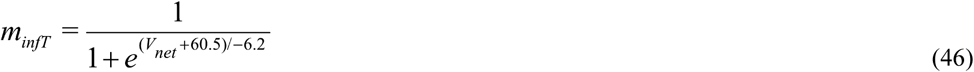

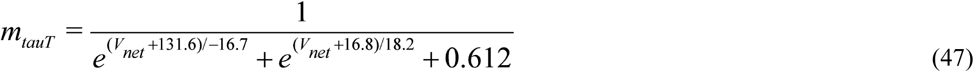

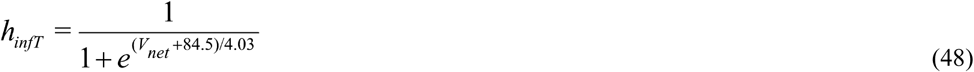

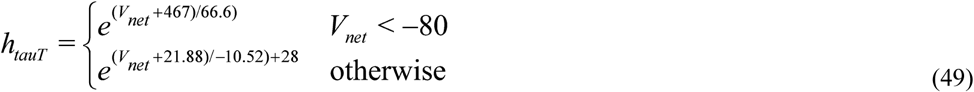

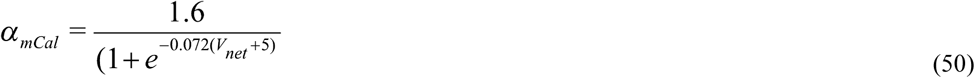

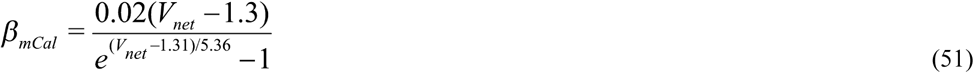

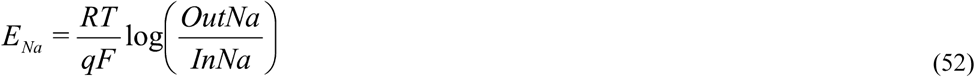

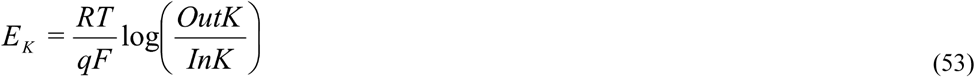

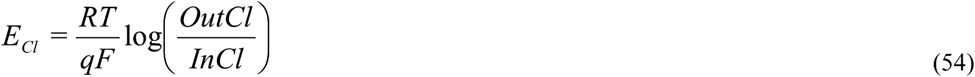

